# Altered spread of waves of activities at large scale is influenced by cortical thickness organization in temporal lobe epilepsy: a MRI-hdEEG study

**DOI:** 10.1101/2023.07.05.547809

**Authors:** Gian Marco Duma, Giovanni Pellegrino, Giovanni Rabuffo, Alberto Danieli, Lisa Antoniazzi, Valerio Vitale, Raffaella Scotto Opipari, Paolo Bonanni, Pierpaolo Sorrentino

## Abstract

Temporal lobe epilepsy (TLE) is a brain network disorder characterized by alterations at both the structural and the functional level. It remains unclear how structure and function are related and whether this has any clinical relevance. In the present work, we adopted a novel methodological approach investigating how network structural features influence the large-scale dynamics. The functional network was defined by the spatio-temporal spreading of aperiodic bursts of activations (neuronal avalanches), as observed utilizing high-density electroencephalography (hdEEG) in TLE patients. The structural network was modeled as the region-based thickness covariance. Loosely speaking, we quantified the similarity of the cortical thickness of any two brain regions, both across groups, and at the individual level, the latter utilizing a novel approach to define the personalized covariance network (pCN). In order to compare the structural and functional networks (at the nodal level), we studied the correlation between the probability that a wave of activity would propagate from a source to a target region, and the similarity of the source region thickness as compared to other target brain regions. Building on the recent evidence that large-waves of activities pathologically spread through the epileptogenic network in TLE, also during resting state, we hypothesize that the structural cortical organization might influence such altered spatio-temporal dynamics. We observed a stable cluster of structure-function correlation in the bilateral limbic areas across subjects, highlighting group specific features for left, right and bilateral TLE. The involvement of contralateral areas was observed in unilateral TLE. We showed that in temporal lobe epilepsy alterations of structural and functional networks pair in the regions where seizures propagate and are linked to disease severity. In this study we leveraged on a well-defined model of neurological disease and pushed forward personalization approaches potentially useful in clinical practice. Finally, the methods developed here could be exploited to investigate the relationship between structure-function networks at subject level in other neurological conditions.

## 1. INTRODUCTION

The human brain is organized along multiple nested layers that span from the microscale (e.g. the molecular level), to the mesoscale (e.g. the cytoarchitectonics), all the way up to whole-brain structures (e.g. the white-matter bundles and the brain geometry) (Hilgetag & Goulas, 2020). Such hierarchical structure induces resonances resulting in spontaneous collective bursts of activity that spread across the brain (Deco et al., 2013). Alterations of large-scale brain dynamics have been detected in multiple neurological diseases (Li et al., 2021; Sorrentino, Rucco, et al., 2021).

This is of particular relevance in epilepsy, which has come to be considered a network disorder affecting not only the local node (i.e., the epileptogenic zone; EZ) but also the whole-brain network functioning (Bartolomei et al., 2017; Gotman, 2008). From the structural standpoint, MRI studies have demonstrated that patients with epilepsy display cortical atrophy not only in the EZ but also more diffusely, supporting the concept of epileptogenic network (Caciagli et al., 2017; Labate et al., 2011). A recent study from the ENIGMA-epilepsy study group has evidenced diffused alteration of the cytoarchitectonic and morphological cortical organization in patients with temporal lobe (TLE) and idiopathic generalized epilepsy (IGE) (Larivière et al., 2022). In line with these results, neuroimaging and electrophysiological studies have shown patterns of altered functional connectivity impacting the whole-brain, both during ictal and interictal activity (Courtiol et al., 2020; Duma et al., 2021; Wirsich et al., 2016). A recent work from our group used resting state electroencephalography (rs-EEG) to investigate the propagation of bursts of activities on the large-scale in patients with TLE (Duma et al., 2023). We showed that, even in the inter-ictal period, in the absence of epileptiform abnormalities, alteration of the propagation of waves of activities on the large-scale clustered onto key brain areas with respect to seizure onset and propagation. However, it is important to understand how functional networks are constrained by structural properties (Van Diessen et al., 2013; Voets et al., 2012).

Recent studies suggest that including regional heterogeneity, for example with respect to morphological, cytoarchitectonics and neuromodulatory information, is fundamental to understand and to model the role of structural organization in constraining the spatio-temporal dynamics (Suarez et al., 2020). The addition of brain morphology in the investigation of the structure-function relationship is particularly relevant in epilepsy since brain structural organization is affected by the pathology, for instance as cortical thinning in the context of focal epilepsy (Galovic et al., 2019). In this study, we hypothesized that cortical thickness distribution in the brain may be linked to the altered spreading of the large-scale perturbations in relation to epilepsy. To test this hypothesis, we used a multimodal dataset made of source-reconstructed high-density electroencephalography (hdEEG) and structural MRI from 59 patients with TLE. The model of TLE is of particular interest because it represents a rather well-defined group of electroclinical conditions with lower clinical heterogeneity compared to other epilepsy forms. Previous studies investigating structural and functional anomalies in epilepsy have intensively relied on this model (Larivière et al., 2020; Song et al., 2022; Xie et al., 2023). Leveraging on our previous work on altered avalanche spreading in TLE (Duma et al., 2023), we expect to identify a relationship between the organization of the cortical morphology and the propagation of the activity bursts, and specifically so in the epileptogenic network. Moreover, in the light of the relationship between thickness and disease duration (Galovic et al., 2019), as well as the exposure to antiseizure medications (Bittigau et al., 2002; Kim et al., 2007), we explored the dependence of structure-function link in relation to the age of onset, the epilepsy duration and the number of antiseizure medications (ASMs).

To characterize the fast bursts of activities at the whole-brain level, we deployed the avalanche transition matrix (ATM), a recently developed analytical tool that captures the probability of any two regions being successively recruited by spontaneous neuronal avalanches (Rucco et al., 2020; Sorrentino, Rucco, et al., 2021; Sorrentino et al., 2022). The ATM provides a measure of relationship across regions, resulting in a functional network. In order to better characterize the link between functional and morphological brain configurations, we adopted a network-level method also for the thickness distribution. To this end we analyzed the structural covariance network (SCN), a measure adopted in the workflow of the ENIGMA (Larivière et al., 2022). The SCN describes the existence of correlated anatomical measurements, such as cortical thickness, between pairs of brain regions, proving a network measure of the cortical thickness organization (Wright, 1999). By providing information on the morphology heterogeneity of the cortex, the SCN can be used to elucidate the relationship between the structural and functional connectome. However, as the SCN has been designed as a group metric, it is not trivial to extract individual information.

In this work, by generalizing the co-fluctuation framework (Sorrentino et al., 2023; Esfahlani et al., 2020) we have been able to obtain a subject-wise measure for the SCN (see Method sections) allowing the investigation of the structure-function relationship in an individualized fashion. The individual level SCN will be from here on referred to as personalized covariance network (pCN).

We studied the structural-functional relationship by comparing structural and functional matrices, first at the group level, and then in a subject-specific fashion. In fact, utilizing the subject-wise investigation, we aimed at capturing fluctuations that may be related with pathology-related variables, which is relevant for the clinical translation of our approach. This is in line with the application of individual network-level measurements together with clinical variables in the framework of personalized diagnosis (Jirsa et al., 2017).

## 2. METHOD

### 2.1 Participants

We retrospectively enrolled 70 patients with temporal lobe epilepsy, who underwent high density electroencephalography (hdEEG) for clinical evaluation in 2018-2021 at the Epilepsy and Clinical Neurophysiology Unit, IRCCS Eugenio Medea cited in Conegliano (Italy). The diagnosis workflow included clinical history and examination, neuropsychological assessment, long term surface Video EEG (32 channels) monitoring, hdEEG resting state recording, 3T brain magnetic resonance imaging, and positron emission tomography (PET) as an adjunctive investigation in selected cases. The diagnosis of temporal lobe epilepsy was established according to the ILAE guidelines. Eleven subjects were excluded due to the poor quality of the MR images resulting in a sample of 59 patients with TLE (29 left-TLE, 17 right-TLE, 13 bitemporal TLE). A description of patients’ demographic and clinical characteristics is provided in Table 1. The study protocol was conducted according to the Declaration of Helsinki and approved by the local ethical committee.

**Tab 1.**
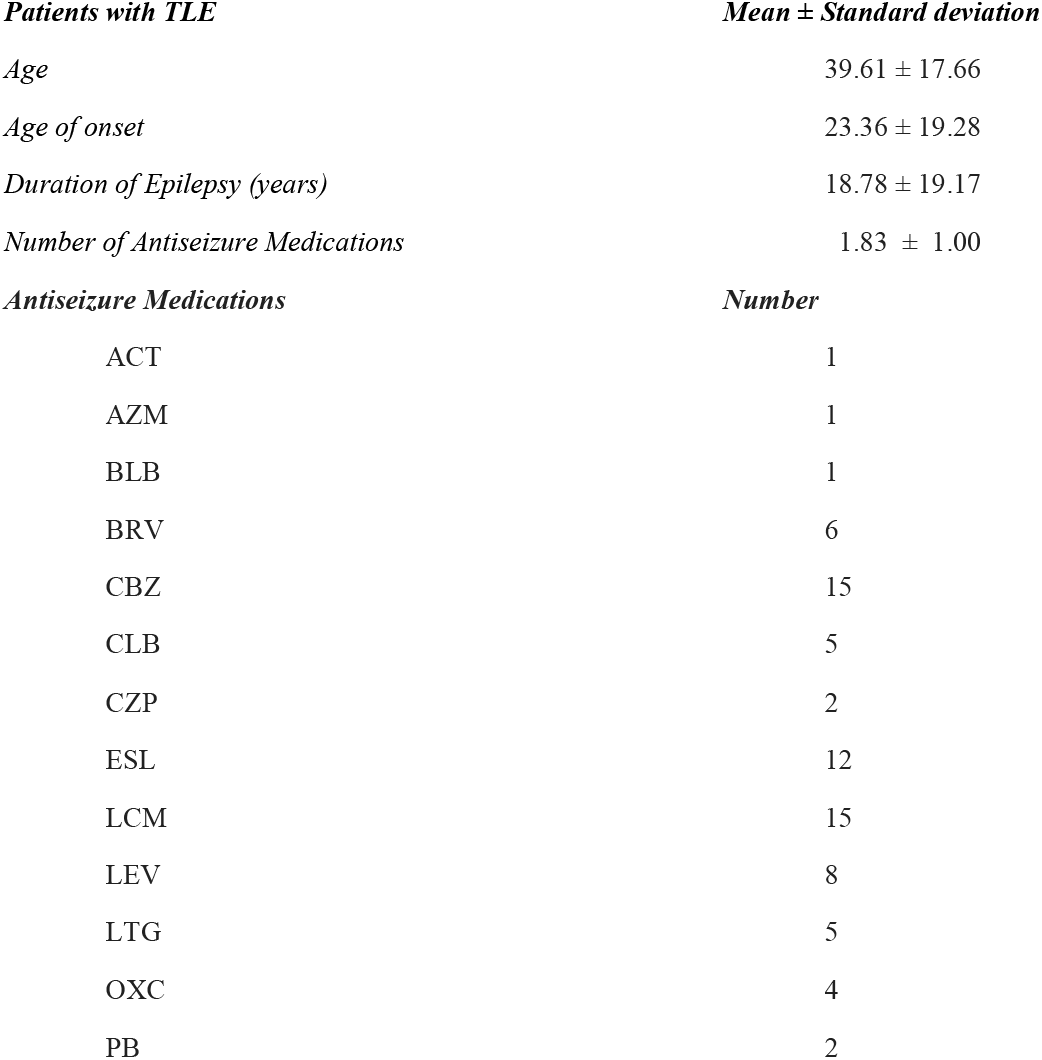
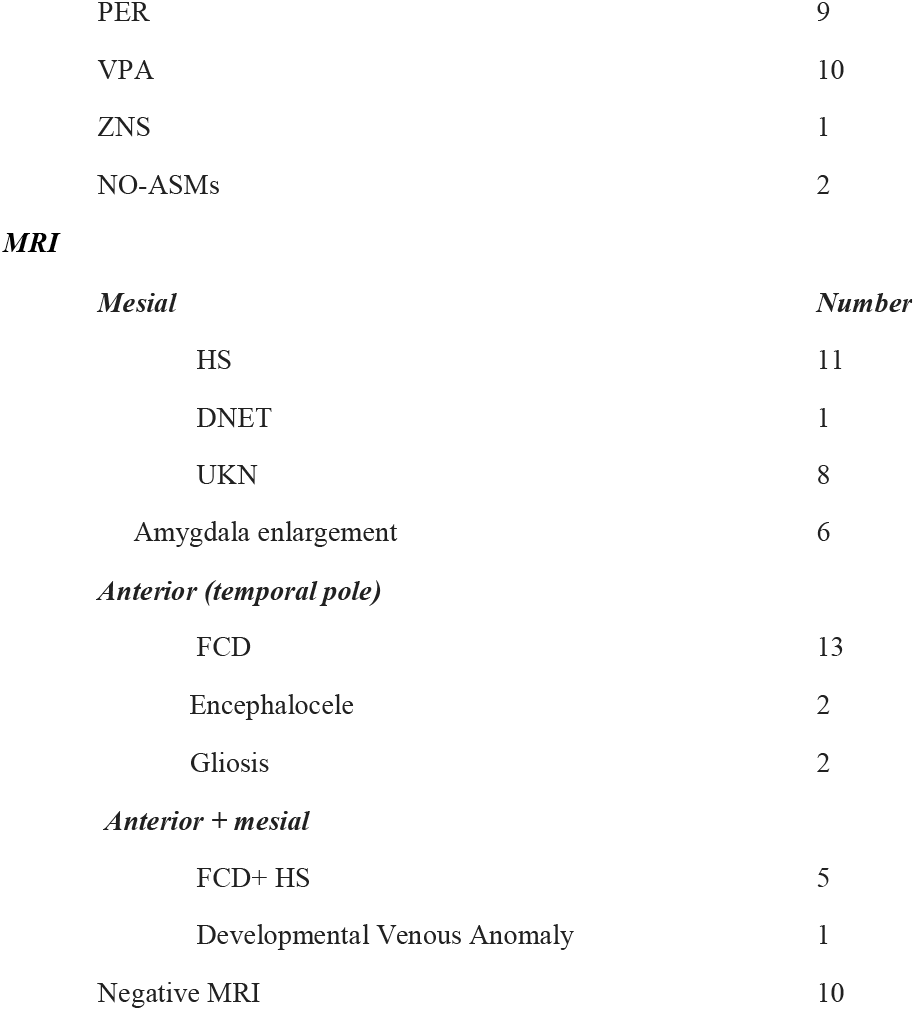
The table describes the demographic and clinical characteristics of the patients with temporal lobe epilepsy, along with the scores of the neuropsychological tests. MRI abnormalities are reported by sublobar localization. The continuous variables are reported as mean ± standard deviation. Antiseizure Medication abbreviations: ACT= acetazolamide, AZM = acetazolamide; BRV = brivaracetam, CBZ = carbamazepine, CLB = clobazam, CZP = clonazepam, ESL= eslicarbazepine, LCM = lacosamide, LEV = levetiracetam, LTG = lamotrigine, OXC = oxcarbazepine; PB = phenobarbital, PER = perampanel, VPA= valproic acid, ZNS = zonisamide, NO-ASMs = no pharmacological treatment. Abbreviation of the identified anomalies on the magnetic resonance imaging: FCD= focal cortical dysplasia, HS=hippocampal sclerosis, DNET = dysembryoplastic neuroepithelial tumors, UKN = unknown.

### 2.2 Morphological measures and covariance networks

We used the individual MRI anatomy in order to generate individualized head models for the patients with TLE. The anatomical MRI for source imaging consisted of a T1w 1 mm isotropic 3D acquisition. The MRI was segmented in skin, skull and gray matter using the segmentation pipeline of the Computational Anatomy Toolbox (CAT12; Gaser et al., 2022). Successively, we computed the cortical thickness using CAT12, which yields a morphological value for each vertex of the brain mesh. We used the Desikan-Killiany parcellation (Desikan et al., 2006) and then by averaging across all the vertices within each region of the atlas, we obtained the morphological index value *Xi* (*s*) for each region *i* of the atlas and for each subject *s*. The structural covariance matrix (SCN) of a group of subjects is computed by estimating the inter-regional Pearson’s correlation across subjects of the cortical index between all possible pairs of regions (Romero-Garcia et al., 208). Each element of the SCN matrix in defined as

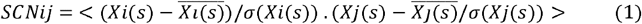

where 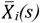 denotes the mean cortical index of region *i* across all subjects, and *σ*(*Xi*(*s*)) denotes its variance. The formula above defines Pearson’s correlation i.e., the product of z-scored cortical indices, averaged across all subjects. Thus, the construction of the SCN relies on the identification of spatial patterns of morphometric similarities between brain regions within a group of subjects. The SCN matrix can be expressed at the single-subject level by defining the personalized covariance network (pCN) as

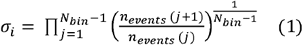

This way the SCN is the result of averaging across individual SCN(s) i.e.,

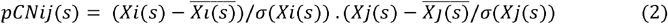

Hence, the pCN can be understood as the contribution of each subject to the group-level correlation. The above deconstruction of the pCN was inspired by a recently introduced edge-centric approach to functional connectivity (Esfahlani et al. 2020, Sorrentino et al., 2023). In doing so, the morphological information was structured as a 3-D matrix 68 (ROIs) × 68 (ROIs) × 60 (subjects), representing all individual pCN(s). This derivation of the pCN allowed to investigate the relationship of the cortical network with the functional organization derived from the avalanche transition matrix (ATM), which had the same 3D structure (see section 2.7) both at the group and individual level.

### 2.3 Resting State EEG recording

The hdEEG recordings were obtained using a 128-channel Micromed system referenced to the vertex. Data was sampled at 1,024 Hz and the impedance was kept below 5kΩ for each sensor. For each participant, we recorded 10 minutes of closed-eyes resting state while comfortably sitting on a chair in a silent room.

### 2.4 EEG pre-processing

Signal preprocessing was performed via EEGLAB 14.1.2b (Delorme & Makeig, 2004). The continuous EEG signal was first downsampled at 250 Hz and then bandpass-filtered (0.1 to 45 Hz) using a Hamming windowed sinc finite impulse response filter (filter order = 8250). The signal was visually inspected to identify interictal epileptiform discharges (IEDs) by GMD, AD and PB and then segmented into 1-sec long epochs. Epochs containing IEDs activity were removed. Epoched data underwent an automated bad-channel and artifact detection algorithm using the TBT plugin implemented in EEGLAB. This algorithm identified the channels that exceeded a differential average amplitude of 250μV and marked those channels for rejection. Channels that were marked as bad in more than 30% of all epochs were excluded. Additionally, epochs having more than 10 bad channels were excluded. We automatically detected possible flat channels with the Trimoutlier EEGLAB plug in within the lower bound of 1μV. We rejected an average of 17.34 ± 11.22 (SD) epochs due to spikes and 5.54 ± 3.82 (SD) due to artifacts. The preprocessing analysis pipeline has been applied by our group in previous studies investigating both task-related and resting state EEG activity (Duma, Di Bono, Mento, 2021; Duma et al., 2023). Data cleaning was performed with independent component analysis (Stone, 2002), using the Infomax algorithm (Bell & Sejnowski, 1995) implemented in EEGLAB. The resulting 40 independent components were visually inspected and those related to eye blinks, eye movements, muscle and cardiac artifacts were discarded. The remaining components were then projected back to the electrode space. Finally, bad channels were reconstructed with the spherical spline interpolation method (Perrin et al., 1989). The data were then re-referenced to the average of all electrodes. At the end of the data preprocessing, each subject had at least 8 minutes of artifact-free signal.

### 2.5 Cortical Source modeling

The resulting individual surfaces from CAT12 were then imported in Brainstorm (Tadel et al., 2011), where three individual surfaces adapted for Boundary Element Models (BEM) were reconstructed (inner skull, outer skull and head) and the cortical mesh was downsampled at 15,002 vertices. The co-registration of the EEG electrodes was performed using Brainstorm, by projecting the EEG sensor positions on the head surface with respect to the fiducial points of the individual MRI. We applied manual correction of the EEG cap on the individual anatomy whenever needed, prior to projecting the electrodes on the individual head surface via Brainstorm. We then derived an EEG forward model using 3-shell BEM model (conductivity: 0.33, 0.165, 0.33 S/m; ratio: 1/20) (Goncalves et al., 2003) estimated using OpenMEEG method implemented in Brainstorm (Gramfort et al., 2010). Finally, we used the weighted minimum norm imaging (Hämäläinen & Ilmoniemi, 1994) as the inverse model, with the Brainstorm’s default parameters setting.

### 2.6 Avalanche estimation

Similarly to our previous work using neuronal avalanches (Duma et al., 2023), we extracted the activity of a total of 68 regions of interest (ROIs) from the Desikan-Killiany atlas (Desikan et al., 2006). The ROIs time series were obtained by averaging the activity across the vertices composing each ROI. To study the dynamics of brain activity, we estimated “neuronal avalanches” from the source-reconstructed ROI time series. Firstly, the time series of each ROI was discretized, by calculating the z-score over time and then setting positive and negative excursions beyond a threshold as 1, and the rest of the signal as 0. A neuronal avalanche begins when, in a sequence of contiguous time bins, at least one ROI is active (i.e., above threshold), and ends when all ROIs are inactive (Beggs & Plenz, 2003; Shriki et al., 2013; Sorrentino, Seguin, et al., 2021). The total number of active ROIs in an avalanche corresponds to its size. These analyses require the time series to be binarized. This is done to ensure that one is capturing critical dynamics, if present. To estimate the suitable time bin length, for each subject, each neuronal avalanches, and each time bin duration, the branching parameter ⍰ was estimated (Haldeman & Beggs, 2005). In fact, systems operating at criticality typically display a branching ratio ∼1. The branching ratio is calculated as the geometrically averaged (over all the time bins) ratio of the number of events (activations) between the subsequent time bin (descendants) and that in the current time bin (ancestors), and then averaging it over all the avalanches (Bak et al., 1987). More specifically:

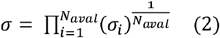

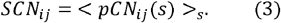

where ⍰_*i*_ is the branching parameter of the *ith* avalanche in the dataset, *N*_*bin*_ is the total amount of bins in the *i-th* avalanche, *n*_*events*_ *(j)* is the total number of events active in the *j-th* bin, and *N*_*aval*_ is the total number of avalanche in the dataset. In our analyses the branching ratio was 1 for bin = 2 (corresponding to bins of 8 ms).

### 2.7 Avalanche transition matrices

An avalanche-specific transition matrix (ATM) was calculated where element (*i, j*) represented the probability that region j was active at time *t +⍰*, given that region *i* was active at time ***t***, where *⍰* ∼8 ms. The ATMs were averaged element-wise across all avalanches per each participant, and finally symmetrized to obtain individualized ATM.

### 2.8 Correlation analysis

At first, we correlated the mean thickness value of the ROIs with their mean transitivity (averaged transitivity value of the ROI from ATM) across subjects. Then, we computed the correlation between the network organization of the brain morphology (i.e., pCN) and the intrinsic functional organization (i.e., ATM). The pCN matrix was structured similarly to the the 3D ATMs matrix, i.e. 68 (ROIs) × 68 (ROIs) × 60 (subjects). We then carried out correlations at the global and subject-wise level. Specifically, we extracted the thickness and avalanche transition value from the 3D matrix for each *i-th* node with the rest of the brain across subjects, yielding a 68 (ROIs) × 60 (subjects) matrix. We vectorized these matrices, and we applied permutation-based (10.000 permutation) Spearman’s correlation, using the max-statistic for the *p*-value correction (Groppe et al., 2011). The result represents the global correlation of the thickness covariance value and the transition probability of a specific cortical (*i-th* node) area across subjects (see Fig. 1C). Furthermore, it is possible to extract ROIs × ROIs slices from the 3D thickness covariance matrix, which represent the pCN. We then used Spearman’s correlation between the subject-level pCN and ATM, obtaining a value between the individual covariance of a specific node and its transition probability (see Fig. 1C). Successively, for each region, we provided the percentage of times for a region showing a significant pCN-ATM relationship across subjects. Finally, we correlated, using Spearman’s correlation, the number of significant regions for each subject with the age of onset, the disease duration and the number of ASMs.

**FIG 1.**
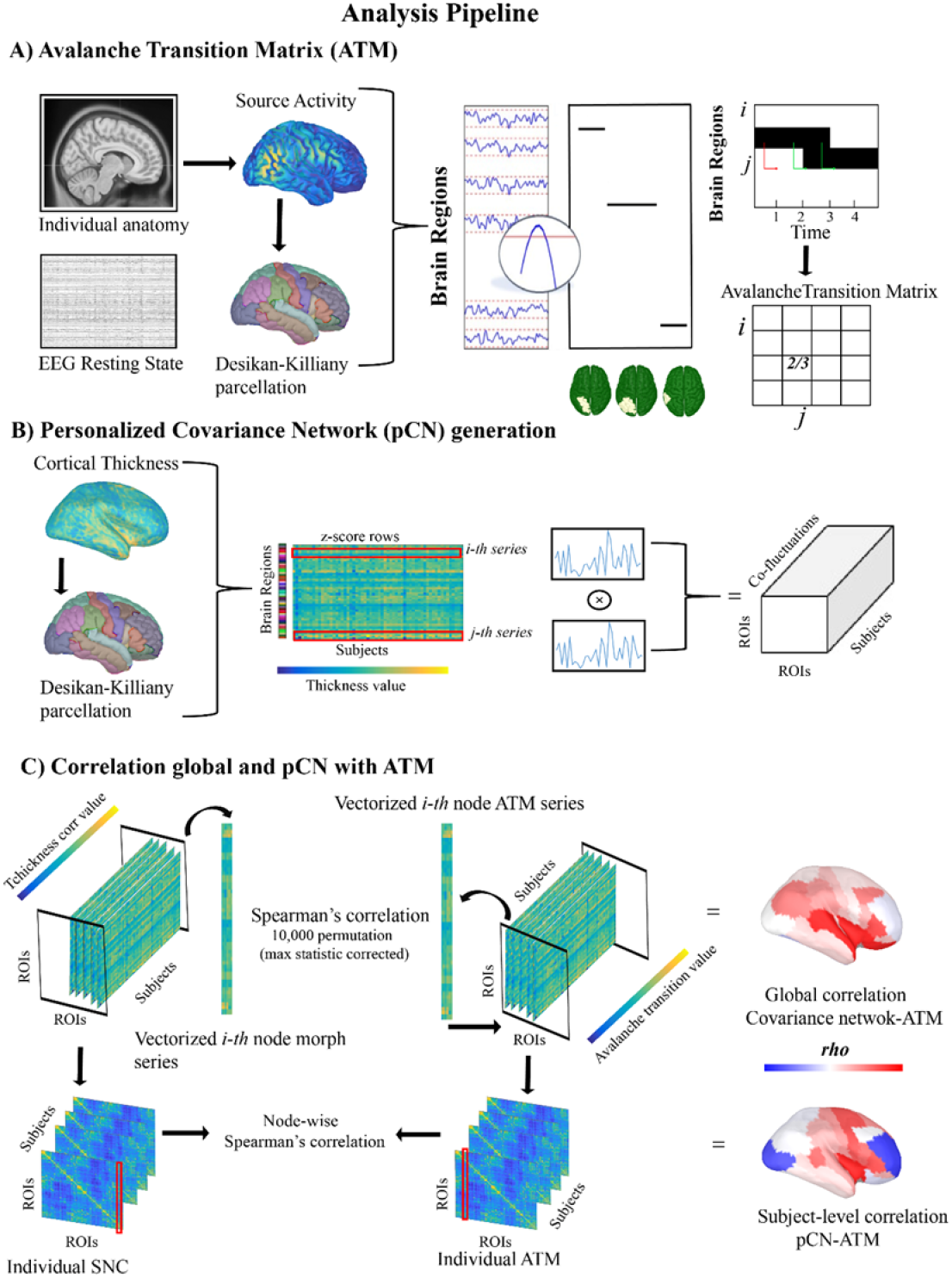
Analysis pipeline. The present figure shows the analytical steps in the analysis process. Panel A displays the pipeline starting from the structural (magnetic resonance imaging) and functional (high density electroencephalography), necessary to obtain the avalanche transition matrix. Panel B shows the computation of the personalized covariance network, from the mean thickness value of the region of interest. Finally, panel C represents the analytical step in the structure-function network correlation both at the group and subject-wise level.

## 3. RESULTS

### 3.1 Group level analysis

For the sake of simplicity, in this section we report the maximum structure-function correlation Sperman’s *rho* for each group. The *rho* and the relative *p*-value for each significant region and each group are provided in the supplementary materials. When considering the correlation between the average cortical thickness of a ROIs and its average transition probability, no significant results were found (*p-val* > .05). Group-wise significant correlation between structure and function was detected when considering the network level both at the structural (thickness co-fluctuation) and functional level (ATMs). Figure 2 shows the unthresholded (panel A) and the corresponding thresholded (panel B) surface maps (*p-val* < .01; max *rho* = .092; for all the regions and their value see Supplementary Tab.1). One can note that the statistical significances displayed a characteristic topography of the regions where structure and function correlated. In particular, structure-function correlation clustered in the temporal regions, including the superior and middle temporal poles, the insula, the anterior middle cingulate and the parahippocampal cortices (Desikan-Killiany atlas based) (see Fig. 2B). The *rho* value was positive, indicating that larger thickness co-fluctuations in these regions correspond to an increased probability of neuronal avalanche spreading through them. We repeated the analysis for the left-TLE, right-TLE and also for bitemporal patients separately. On the one hand, our findings highlight a significant contribution of the temporal areas contralateral to the epileptogenic zone. In the left-TLE we observed a significant correlation (*p-val* < .01; max *rho* = .107; for all the regions and their value see Supplementary Tab.2) between the thickness co-fluctuation and the ATM, mainly involving the right temporal areas, i.e. the banks of the superior temporal sulcus and the middle temporal lobe, as well as the parietal (supramarginal and superior parietal) and the bilateral frontal areas (rostral middle frontal and pars triangularis) (see Fig. 2C-D). On the other hand, right-TLE patients are characterized by the involvement of left middle temporal area, as well as the bilateral insular areas in conjunction with caudal and middle frontal regions (*p-val* < .01; max *rho* = .145; see Fig. 2E-F; for all the regions and their value see Supplementary Tab.3). Finally, the bitemporal group shows a correlation of the bilateral banks of the superior temporal sulcus and the insula, as well as of the superior temporal, together with the left precentral and right anterior cingulate (*p-val* < .01; max *rho* = .151; see Fig. 2G-H; for all the regions and their value see Supplementary Tab.4).

**FIG 2.**
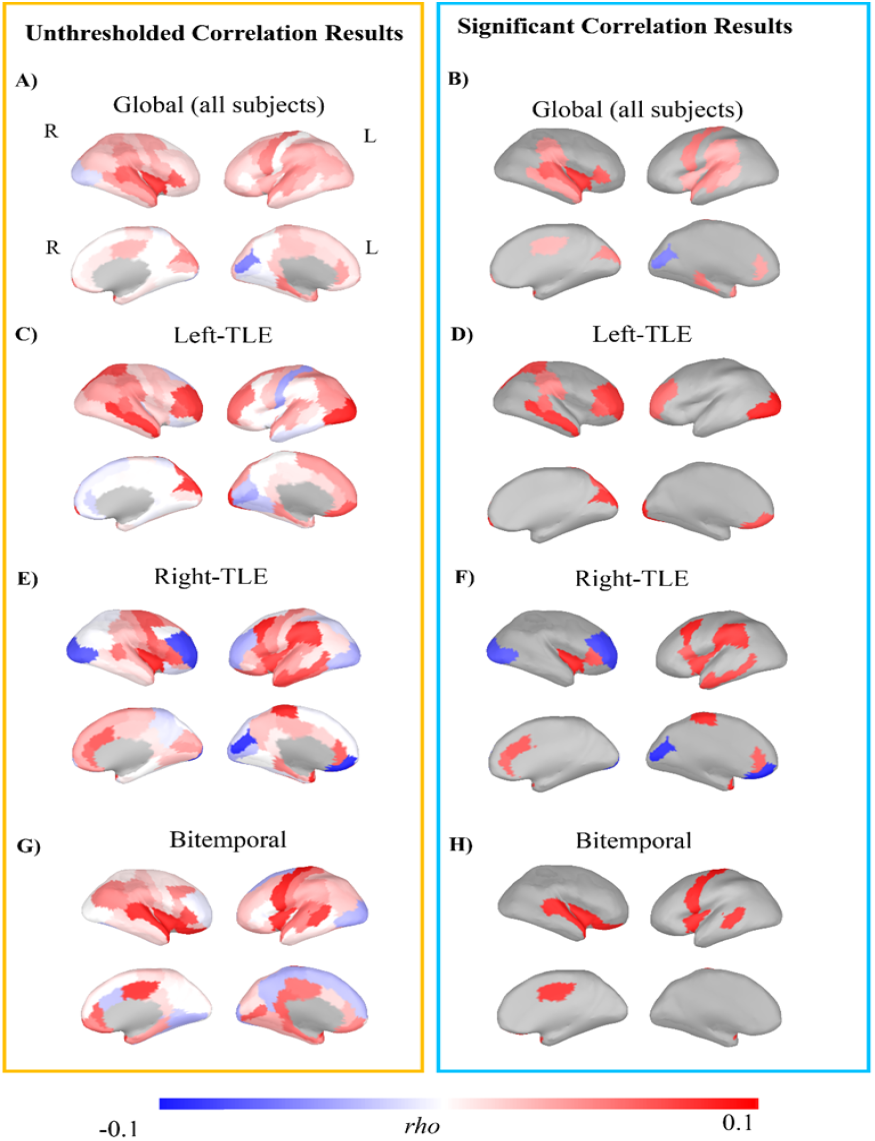
Group level structure-function networks correlation. The present figure displays on the left panels the untresholded correlational results at the group level between the covariance network and the avalanche transition matrix. By contrast on the right panel are shown the thresholded (*p* < .01) results after false discovery rate correction. Specifically, panel B shows the results taking into account the full sample of 60 subjects. Additionally, panels D, F and H show the group level results for patients with left and right unilateral temporal lobe epilepsy, and bilateral temporal lobe epilepsy.

### 3.2 Subject level results

Subject level results are provided as examples for two randomly-selected cases, whereas the maps of all the participants are available online at the following link of the Open Science Framework (https://osf.io/9zj3v/?view_only=94a3287ed35e4040aa791076cc032190). One of the patients reported in the manuscript (SJ-38) is a right TLE patient showing a well-defined pCN-ATM relationship with involvement of the temporal areas contralateral to the epileptogenic site (max *rho* = .448; *p-val* < .01; see Fig. 3-B; for all the regions and their value see Supplementary Tab.5). The second subject that we report (SJ-41) showed a distributed structure-function relationship in bilateral temporal areas, including parahippocampal cortex, superior and inferior temporal, as well as frontal areas (max *rho* = .413; *p-val* < .01; see Fig. 3-B; for all the regions and their value see Supplementary Tab.6). As a summary of the subject-wise analysis, we provide the percentage of times across subjects in which a region displays a significant structure-function relationship by region. The temporal and fronto-central areas, as well as the insular and precentral gyrus, the temporal parietal junction and the cingulate cortex (see Fig. 3 A) are the most consistent area with a larger probability of aperiodic bursts propagation in relation to the morphological configuration. structure-function relationship.

**FIG 3.**
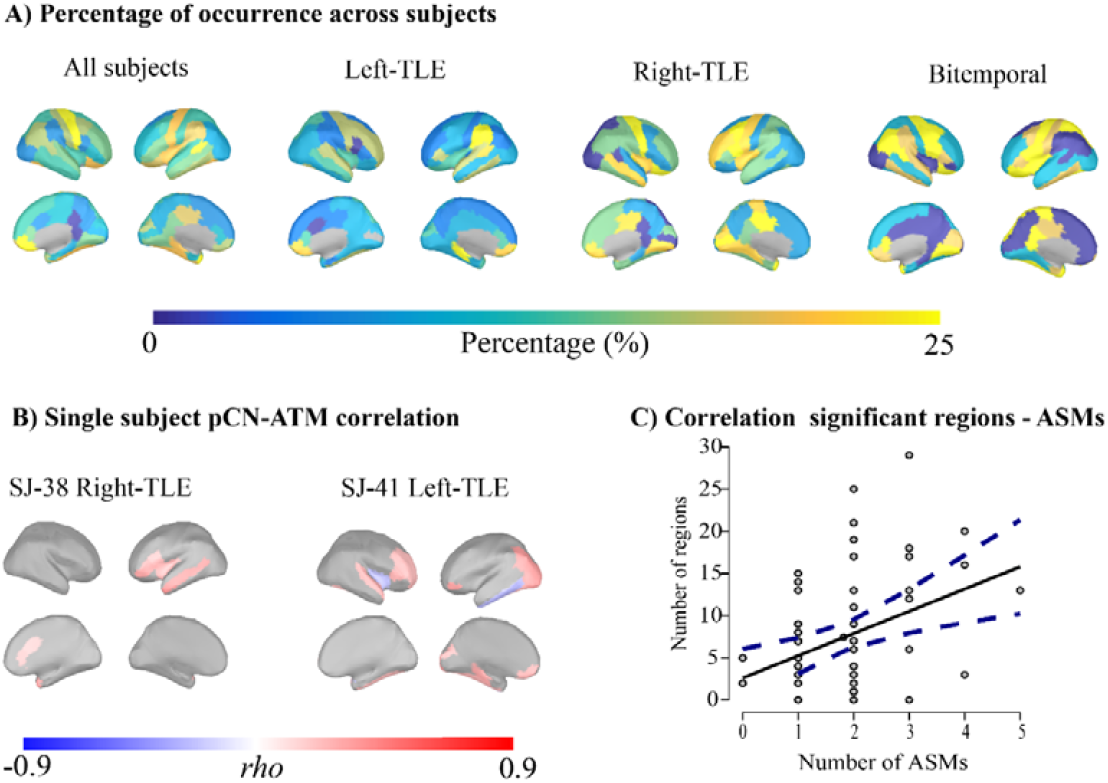
Additional results. In the Panel A of the figure are represented, as a summary of the individual level statistic, the percentage of time across subjects in which a region shows a significant structure-function relationship. Panel B shows the pCN-ATM correlation of two subjects randomly selected. Finally, the correlation between the number of regions with a significant pCN-ATM correlation and the number of antiseizure medications is displayed in panel C. The dashed blue lines represent the confidence intervals of 95%.

### 3.3 Relationship with clinical variables

A significant relationship between the number of regions with a significant structure-function link and the number of antiseizure medications was found (*rho =* .28, *p =* .030) (see Fig. 3 C). By contrast, no significant correlations (*p* > .05) were found with the age of onset nor with epilepsy duration.

## 4 DISCUSSION

In the present work we investigated the relationship between cortical morphology and brain activity in TLE. We focused on cortical thickness as a marker of brain morphology, as this is altered in TLE (Bernhardt et al., 2010, 2011; Caciagli et al., 2017; Larivière et al., 2022). As for brain activity, we focused on the spontaneous spatio-temporal dynamics from hdEEG data. Specifically, we computed the avalanche transition matrices (ATM), that is a mathematical tool accommodating the non-linearities of the large-scale brain dynamics and the corresponding multimodal dynamics that they generate (Duma et al., 2023; Rucco et al., 2020; Sorrentino et al., 2022; Sorrentino, Seguin, et al., 2021). We confirmed our hypothesis that the avalanche spread more across regions that are more similar in their structural features, namely the network organization of cortical thickness. Larger covariation of thickness cortex facilitates/enhances avalanche propagation. These results expands previous knowledge that activity synchronization and propagation depends upon cortical thickness in healthy individuals (Schuler et al., 2022) by adding the information coming from the non-linearities of the large-scale brain dynamics. It is interesting to note that the relationship between cortical thickness and avalanche spreading involves the limbic system and the regions where temporal lobe seizures are known to propagate (Jo et al., 2019; Kubota et al., 2013; Yoo et al., 2014), such as the bilateral temporal areas, the posterior temporal regions, the insula, the parahippocampal and the cingulate cortex (see Fig 2B). Interestingly, these are also regions whose cortical thickness has been found altered in previous studies, possibly as an effect of the recurrent spread of seizure activity (Abdelnour et al., 2015; Bonilha et al., 2010; Galovic et al., 2019). Importantly, the structure-function relationship is not observed when the thickness of a region is directly related to the mean avalanche transitivity, but only when the network level is taken into account. These findings imply that the morphology of a single region is not sufficient to explain the spreading of the aperiodic activity by itself. In fact, neuronal avalanches are a product of large-scale brain activity, meaning that the transitivity in one region is highly dependent on the network activity. For this reason, a whole-brain structural configuration perspective can capture the relationship between neuronal large scale dynamics and brain structure. This is in line with the recent findings that cortical activity can be better understood as resulting from excitations of fundamental, resonant modes of the brain’s structure (Pang et al., 2023). In fact, the pCN considers the covariance of one region with the others across subjects, or intra-individually, partly capturing the geometric configuration of the whole-cortex. Our findings suggest that the relationship between structure and activity is significant in both temporal lobes in patients with TLE. To investigate the structure-function relationships in different TLE subgroups, we then performed additional analysis by dividing our clinical population into left, right and bilateral TLE. This comes with the price of a smaller sample size for each group. This analysis revealed that patients with bilateral TLE show a significant bilateral temporal thickness-avalanches relationship (see Fig. 2H) Surprisingly, unilateral TLE was associated with a stronger structure-function link in the contralateral temporal lobe, together with contralateral temporo-parietal junction and bilateral prefrontal areas (see Fig. 2D and F). The thinning of the contralateral temporal areas has been described in unilateral temporal lobe epilepsy with a greater involvement of the right-TLE (Park et al., 2022; Seidenberg et al., 2005), but to a lesser degree as compared to the ipsilateral cortex. One possibility is therefore that the contralateral temporal lobe is less affected by cortical thinning, resulting in an enhancement of the avalanche spread. One alternative explanation is that the contralateral compensatory plastic changes occur in TLE and explain the increased connectivity in the contralateral regions (Bettus et al., 2009). Overall, our results align well with the interpretation of epilepsy as a network disorder. While previous studies have highlighted either impairment of structure or functional data, here we demonstrate how these two properties closely interact, providing a comprehensive view leveraging on multimodal data. We further tried to account for variability in our sample by pushing forward an attempt to bring the study of structure-function relationship at the subject level, acknowledging that even within homogeneous TLE groups each subject has their own specificities (see Fig. 3B). Subject-level analysis confirmed and strengthened group-level results. The link between thickness and avalanche spreading was stronger at the individual level, suggesting that the group-level analysis may underestimate the relationship due to inter-individual differences. Some cortical regions were consistently recruited by ongoing avalanches, particularly in the brain regions that are structurally altered in TLE (Fig 3A). Finally, with regards to the possible relation between anatomofunctional architecture and clinical variables (age of epilepsy onset, epilepsy duration and number of ASMs) we only found an association between the number of regions with significant structure-function correlation and ASMs, with the former increased in individuals with higher ASMs load. The ASMs load may reflect epilepsy severity, which in turn has been associated to a more diffuse spreading of epileptic activity within the brain (Andrews et al., 2019). Moreover, our previous findings highlighted that patients with TLE are characterized by an iper-integration of the functional networks (Duma et al., 2022). In light of this, we may speculate that a more severe clinical picture, likely requiring a larger number of ASMs, is linked to a widespread dysregulation of both the functional and structural networks, resulting in a less segregated and localized structure-function links. To sum up, in the present work we merge structural and functional imaging in TLE patients, and demonstrate, subject-wise, a relationship between the alterations of the aperiodic dynamics and cortical organization. We provided a methodological insight with a new way to compute a personalized structural covariance, together with an innovative approach investigating the aperiodic brain dynamics, namely the neuronal avalanches. Our approach finds its rationale in the idea that the altered activities in epilepsy might be the result of local structural alterations and the way they affect the resonances that are generated at the whole-brain level. As such, the integration of structural and whole-brain functional data is indispensable. In fact, our results may have clinical relevance for the diagnostic process, especially in terms of the identification of the epileptogenic network (EN). Our findings could represent a first step in the inclusion of the morphological regions heterogeneity to increase the accuracy of the models of brain dynamics in epilepsy (Hashemi et al., 2020; Jirsa et al., 2017; Proix et al., 2018), as suggested by Suarez and colleagues (Suárez et al., 2020). In fact, the inclusion of the morphological configuration to predict the behavior of the brain networks is useful for the surgical treatment in focal epilepsy, as well as to enhance personalized modeling for the optimization of drug delivery or neuromodulatory approaches (Sobayo & Mogul, 2016).

## CONCLUSIONS

In the present work we deployed a novel methodological approach to test the hypothesis that large-scale dynamics is influenced by structural features of the cortex. We observed a stable cluster of correlation in the bilateral temporal and limbic areas across subjects, highlighting group-specific features for left, right and bilateral TLE patients. We developed strategies to bring the investigation to the individual level, confirming group-wise findings and expanding them to the single subject level. We confirmed that TLE is characterized by structural cortical alterations that are intimately related to the alteration of the fast whole-brain functional dynamics. Finally, we showed that the structure-function link has a tight relationship with clinical features such as disease severity. In this study, we leveraged on a well-defined model of neurological disease and pushed forward personalization approaches potentially useful in clinical practice for TLE. Nevertheless, the present methodology may have clinical implications in conditions with a much broader heterogeneity of structural alterations across the brain (e.g., stroke, multiple sclerosis).

## Supporting information

Supplementary Results

## ACKNOWLEDGMENTS

This work was supported by a 2019 “5XMille” to PB, Ricerca Corrente 2023 to GMD funds for biomedical research of The Italian Health Ministry, and by the European Union’s Horizon 2020 research and 569 innovation program under grant agreement No. 945539 (SGA3).

## CONFLICT OF INTEREST

The authors declare no conflict of interest.

## DATA AVAILABILITY STATEMENT

The data that support the findings of this study are available on request from the corresponding author. The data are not publicly available due to privacy or ethical restrictions.

## Notes

### Competing Interest Statement

The authors have declared no competing interest.

### Summary of Updates

We realized that the title was not formally correct, and we changed in a more appropriate one

