## Supplementary Results for "Altered spread of waves of activities at large scale is influenced by cortical thickness organization in temporal lobe epilepsy: a MRI-hdEEG study"

**SUPPLEMENTARY MATERIALS**

| **Regions** | ***rho*** | ***p*-val** |
|---|---|---|
| bankssts L | 0.0427 | 0.0072 |
| bankssts R | 0.0781 | 1.00e-6 |
| cuneus L | -0.0642 | 0.0004 |
| cuneus R | 0.0630 | 1.00e-6 |
| frontalpole R | 0.0546 | 1.00e-6 |
| insula L | 0.0438 | 0.0048 |
| insula R | 0.0922 | 1.00e-6 |
| middletemporal R | 0.0486 | 0.0020 |
| parahippocampal L | 0.0607 | 1.00e-6 |
| parstriangularis R | 0.0768 | 1.00e-6 |
| posteriorcingulate R | 0.0447 | 0.0060 |
| precentral L | 0.0597 | 1.00e-6 |
| rostralanteriorcingulate L | 0.0498 | 0.0016 |
| superiortemporal L | 0.0553 | 0.0004 |
| superiortemporal R | 0.0726 | 1.00e-6 |
| supramarginal L | 0.0450 | 0.0044 |
| supramarginal R | 0.0485 | 0.0036 |
| temporalpole L | 0.0555 | 1.00e-6 |

Supplementary Tab.1. Significant regions with the corresponding Spearman's *rho* and *p*-value for the whole-group analysis.

| **Regions** | ***rho*** | ***p*-val** |
|---|---|---|
| bankssts R | 0.0725 | 0.0008 |
| cuneus R | 0.0994 | 1.00e-6 |
| frontalpole L | 0.0771 | 0.0004 |
| frontalpole R | 0.0957 | 0.0003 |
| lateraloccipital L | 0.1071 | 1.00e-6 |
| medialorbitofrontal L | 0.0806 | 1.00e-6 |
| middletemporal R | 0.0968 | 1.00e-6 |
| parstriangularis R | 0.0763 | 0.0016 |
| rostralmiddlefrontal L | 0.0669 | 0.0028 |
| rostralmiddlefrontal R | 0.0887 | 0.0008 |
| superiorparietal R | 0.0818 | 1.00e-6 |
| supramarginal R | 0.0557 | 0.0096 |

Supplementary Tab. 2. Significant regions with the corresponding Spearman's *rho* and *p*-value for the left-TLE group analysis.

| Regions | *rho* | *p*-val |
|---|---|---|
| caudalanteriorcingulate R | 0.0861 | 0.0028 |
| caudalmiddlefrontal L | 0.0983 | 1.00e-6 |
| cuneus L | -0.1526 | 1.00e-6 |
| insula L | 0.0902 | 0.0012 |
| insula R | 0.1454 | 1.00e-6 |
| lateraloccipital R | -0.0773 | 0.0080 |
| medialorbitofrontal L | -0.1205 | 1.00e-6 |
| middletemporal L | 0.0847 | 0.0024 |
| paracentral L | 0.1273 | 1.00e-6 |
| parsopercularis L | 0.0781 | 0.0076 |
| parstriangularis R | 0.0790 | 0.0048 |
| rostralanteriorcingulate L | 0.0806 | 0.0032 |
| rostralanteriorcingulate R | 0.0798 | 0.0048 |
| rostralmiddlefrontal R | -0.0856 | 0.0020 |
| supramarginal L | 0.0948 | 0.0004 |
| temporalpole L | 0.1033 | 0.0004 |

Supplementary Tab.3. Significant regions with the corresponding Spearman's *rho* and *p*-value for the right-TLE group analysis.

| Regions | *rho* | *p*-val |
|---|---|---|
| bankssts L | 0.0987 | 0.0012 |
| bankssts R | 0.1009 | 0.0020 |
| insula L | 0.1055 | 0.0012 |
| insula R | 0.1359 | 1.00e-6 |
| lateralorbitofrontal R | 0.1087 | 0.0016 |
| posteriorcingulate R | 0.1268 | 1.00e-6 |
| precentral L | 0.1160 | 0.0012 |
| superiortemporal R | 0.1518 | 1.00e-6 |

Supplementary Tab.4. Significant regions with the corresponding Spearman's *rho* and *p*-value for the bilateral TLE group analysis.

| Regions | *rho* | *p*-val |
|---|---|---|
| insula L | 0.3282 | 0.0066 |
| middletemporal L | 0.4483 | 0.0001 |
| parsopercularis L | 0.3142 | 0.0095 |
| parsorbitalis L | 0.3862 | 0.0012 |
| parstriangularis L | 0.3724 | 0.0019 |
| rostralanteriorcingulate R | 0.3230 | 0.0076 |
| temporalpole R | 0.3847 | 0.0013 |

Supplementary Tab.5. Significant regions with the corresponding Spearman's *rho* and *p*-value for the individual level analysis example subject-38.

| **Regions** | ***rho*** | ***p*-val** |
|---|---|---|
| fusiform L | 0.3181 | 0.0086 |
| inferiorparietal L | 0.4133 | 0.0005 |
| lateraloccipital L | 0.3972 | 0.0008 |
| medialorbitofrontal L | 0.3304 | 0.0063 |
| parahippocampal L | 0.3800 | 0.0015 |
| parsorbitalis L | 0.4031 | 0.0007 |
| parstriangularis R | 0.3315 | 0.0061 |
| superiortemporal R | 0.3262 | 0.0070 |

Supplementary Tab.6. Significant regions with the corresponding Spearman's *rho* and *p*-value for the individual level analysis example subject-41.
